# Ethylene Modulates the Phenylpropanoid Pathway by Enhancing *VvMYB14* Expression via the ERF5-Melatonin-ERF104 Pathway in Grape Seeds

**DOI:** 10.1101/2024.09.10.612321

**Authors:** Shiwei Gao, Fei Wang, Shengnan Wang, Shuxia Lan, Yujiao Xu, Xinning Lyu, Hui Kang, Yuxin Yao

## Abstract

Ethylene plays a crucial role in regulating polyphenol metabolism, however the underlying mechanism remains largely unknown. This work demonstrated that ethylene release occurred earlier than melatonin during seed ripening. Ethylene treatment increased the *VvASMT* expression and melatonin content. *VvERF5* was elucidated to bind to the ERE element in the *VvASMT* promoter. *VvERF5* overexpression increased ASMT expression and melatonin content while its suppression generated the opposite results in grape seeds, calli and/or Arabidopsis seeds. Using the promoter of *VvMYB14*, which was strongly induced by melatonin, a melatonin responsive element (MTRE) was identified. *VvERF104* was revealed not only to be strongly induced by melatonin but to bind to the MTRE of the *VvMYB14*. *VvERF104* overexpression and suppression largely increased and decreased the MYB14 expression, respectively, in grape seeds, calli and/or Arabidopsis seeds. *VvMYB14* overexpression widely modified the expression of genes in phenylpropanoid pathway and phenolic compound content in grape seeds. DAP-seq revealed that the MEME-1 motif was the most likely binding sites of *VvMYB14*. *VvPAL*, *VvC4H* and *VvCHS* were verified to be the target genes of *VvMYB14*. Additionally, the roles of *VvERF5*, *VvASMT* and *VvERF104* in mediating ethylene-induced changes in phenylpropanoid pathway were elucidated using their suppressing seeds. Collectively, ethylene increased the *VvMYB14* expression via the pathway of ERF5-melatonin-ERF104 and thereby modified phenylpropanoid pathway.

## Introduction

Ethylene is a gaseous phytohormone that promotes fruit ripening and senescence in climacteric fruits (Bakshi et al., 2015). In addition, ethylene has a crucial regulatory role in nonclimacteric fruits, such as grapes and cherries (Serradilla et al., 2019; Wang et al., 2022). Polyphenols are secondary metabolites that affect fruit quality and antioxidant capacity, and their metabolism is regulated by fruit development (Sreekumar et al., 2014). Ethylene regulates polyphenol metabolism. For instance, treatments with ethylene or its inhibitors affect the composition and content of phenolic compounds in grape berries (Becatti et al., 2014; Xu et al., 2017). Increased levels of total phenolics and p-coumaric acid have been observed in ethylene-treated banana fruits (Tallapally et al., 2020). Ethylene application can also effectively reduce lignin content while increasing the secondary metabolites in ramie (Jie et al., 2023). Ethylene response transcription factors (ERFs) direct specific responses to ethylene signaling by binding directly to promoter regions of ethylene-responsive genes, thereby regulating their expression (Pirrello et al., 2012). In addition, ERFs play a crucial role in regulating fruit ripening, including polyphenol metabolism (Liu et al., 2020). For example, *SmERF1L1* regulates the biosynthesis of tanshinones and phenolic acids in *Salvia miltiorrhiza* (Huang et al., 2019), and *VvERF104* activates the expression of *VvMYBPA2*, enhancing proanthocyanidin synthesis in grape seeds (Zhang et al., 2023). Ethylene also interacts with other hormones, including GA, ABA, and IAA (Arc et al., 2013; Yue et al., 2019; Leubner et al., 2020). ERFs, such as tomato *SlERF.B3* and pear *PuERF2*, also participate in this interaction (Yue et al., 2019; Leubner et al., 2020). However, the interaction between ethylene and melatonin as well as the key transcription factors that integrate ethylene and melatonin signaling remain largely unexplored.

Melatonin is an indolic compound derived from serotonin (5-hydroxytryptamine), and its synthesis in plants is catalyzed mainly by six enzymes, namely tryptophan decarboxylase (TDC), tryptophan hydroxylase (TPH), tryptamine 5-hydroxylase (T5H), serotonin N-acetyltransferase (SNAT), N-acetylserotonin methyltransferase (ASMT), and caffeic acid O-methyltransferase (COMT). Among these, ASMT/COMT and SNAT directly catalyze melatonin synthesis (Back et al., 2016). Melatonin functions not only as a potent antioxidant but also as a multifunctional signaling molecule (Arnao and Hernández-Ruiz, 2019). Melatonin treatment has been shown to alter the secondary metabolite profile of grape skins (Ma et al., 2021), induce proanthocyanidin synthesis in grape seeds (Zhang et al., 2023), and increase the levels of phenolic compounds in goji berries (Wang et al., 2023). Although melatonin signaling in plants remains largely unexplored, it was reported to interact with other signaling molecules, including H_2_O_2_, NO, and various hormones (Caspi et al., 2023). In particular, melatonin promoted grape berry ripening by increasing the levels of ABA, H_2_O_2_, and ethylene (Xu et al., 2018). Plant cell signaling is partially dependent on transcriptional regulatory networks, which consist of circuits of transcription factors and regulatory DNA elements that control the expression of target genes (Priest et al., 2009). Thus, identifying key transcription factors and DNA elements involved in melatonin-regulated processes can provide insights into the underlying mechanisms.

MYB14 is a crucial transcription factor that regulates the phenylpropanoid pathway. In grapes *(Vitis vinifera)*, VvMYB14 regulates stilbene biosynthesis by transactivating *VvSTS* expression (Höll et al., 2013; Mu et al., 2023). In addition, MYB14 regulates other secondary metabolites derived from the phenylpropanoid pathway. For instance, *VvMYB14* affects the accumulation of secondary metabolites in grape calli (Ma et al., 2021) and enhances proanthocyanidin biosynthesis by upregulating the expression of *VvMYBPA1* and *VvMYBPA2* in grape seeds (Zhang et al., 2023). Overexpression of *MtMYB14* was reported to strongly induce proanthocyanidin and mucilage biosynthesis in the seeds and hairy roots of *Medicago truncatula* (Liu et al., 2014). Moreover, MYB14 could increase condensed tannin levels in *Trifolium repens* (Roldan et al., 2019). These findings suggest that MYB14 regulates multiple reactions within the phenylpropanoid pathway. Identifying the specific compounds and target genes regulated by MYB14 will contribute to a better understanding of its functions. Furthermore, MYB14 is known to be involved in melatonin signaling because it is strongly induced by melatonin (Zhang et al., 2023). However, the signal cascade from melatonin to MYB14 remains unclear.

Grape berries are rich in phenolic compounds, including flavonoids such as anthocyanins and proanthocyanidins, as well as nonflavonoid compounds such as phenolic acids and stilbenes (Yilmaz et al., 2015). These phenolics not only substantially affect the flavor of red wine but also act as potent antioxidants, offering potential health benefits. Approximately 30% of the total phenolics in grapes are stored in the seeds (Hanlin et al., 2011), making grape seeds an ideal tissue for studying phenolic compound metabolism. Phenolics are primarily synthesized through the phenylpropanoid pathway. MYB14 regulates the phenylpropanoid pathway; however, its specific target genes are yet to be identified. In addition, the mechanisms through which ethylene and melatonin regulate the phenylpropanoid pathway via MYB14 remain unclear. The present study investigated the function of *VvMYB14* in regulating the phenylpropanoid pathway by identifying its target genes in grape seeds and elucidated the ERF5-melatonin-ERF104 pathway, which mediates ethylene-induced expression of VvMYB14. These findings provide new insights into the molecular mechanisms underlying the role of ethylene signaling in the regulation of melatonin and phenylpropanoid biosynthesis in grape seeds.

## Materials and methods

### Plant materials and growth conditions

Clusters of Merlot grapevines (*Vitis vinifera*), cultivated in an experimental vineyard in Tai-An City, Shandong Province, China, were used as experimental materials. At 67 days after bloom (DAB), grape clusters were treated with a solution of 250 mg·L^−1^ ethephon containing 0.05% Triton X-100 and 10 μL·L^−1^ 1-MCP. The clusters treated with 0.05% Triton X-100 alone served as the control group. In each treatment, the grape clusters were fully immersed in the respective solutions for 30 s, with three biological replicates for each treatment, and each replicate consisting of 30 clusters from five vines.

To induce nonembryogenic calli, seeds from Jianhong grapes were cultured on MS medium supplemented with 0.12 mg·L^−1^ IBA and 1.2 mg·L^−1^ TDZ (Gao et al., 2022). The resulting calli were then subcultured on MS medium containing 30 g·L^−1^ sucrose, 0.60 g·L^−1^ 2-(N-morpholino) ethanesulfonic acid, 8 mg·L^−1^ picloram, 2.5 mg·L^−1^ thidiazuron (TDZ), and 7 g·L^−1^ agar (Zhang et al., 2023). The calli were maintained in a growth chamber at 25°C under a 16-h light/8-h dark photoperiod.

### Determination of the ethylene production rate and melatonin content

The rate of ethylene production was measured using a GC-9A gas chromatograph (Shimadzu, Kyoto, Japan). Three grams of seeds were placed in a 100-mL jar and incubated for 3 h. After incubation, 5 mL of headspace gas was withdrawn using a gas-tight syringe for ethylene production analysis (Ma et al., 2021). The ethylene production rate was calculated based on the ethylene concentration, seed weight, and incubation time (Xu et al., 2018).

Melatonin was extracted and quantified following a modified version of a previously reported method (Xu et al., 2019). Briefly, 1 g of seeds were sonicated in methanol for 20 min to extract melatonin. The supernatant was then centrifuged and evaporated to dryness at 30°C. The resulting residue was dissolved in methanol and purified using a C18 solid-phase extraction cartridge (ProElut; Dikma, China). Melatonin detection was performed using an ACQUITY UHPLC system coupled with a QTof-micro mass spectrometer (Waters, Milford, MA, USA). The UHPLC and MS conditions were identical to those used in our previous study (Ma et al., 2021). Quantification of melatonin was achieved using an external calibration curve based on a melatonin standard.

### Yeast one-hybrid assays, electrophoretic mobility shift assays, and dual-luciferase assay

Yeast one-hybrid (Y1H) assays were performed following the protocol from our previous study (Zhang et al., 2023). The coding sequences (CDS) of *VvERF5*, *VvERF104*, and *VvMYB14* were cloned into the pGADT7 vector, whereas promoter fragments from the target genes were cloned into the pHIS2 vector. These resulting plasmids were transformed into the yeast strain Y187 and plated on SD/-Trp/-Leu medium. The Y1H assay was conducted using a Y1H Library Screening Kit (Clontech, Mountain View, CA, USA) according to the manufacturer’s instructions.

For electrophoretic mobility shift assays (EMSAs), recombinant HIS-tagged *VvERF5*, *VvERF104*, and *VvMYB14* proteins were expressed using the pET-32a vector and purified with a HIS-tag purification column (Beyotime, Shanghai, China). DNA probes containing the ethylene response element (ERE), melatonin response element (MTRE), or MEME-1 element were synthesized and labeled with biotin. EMSAs were performed according to the protocol provided in the LightShift Chemiluminescent EMSA Kit (Beyotime, Shanghai, China).

For the luciferase (LUC) assay, promoter fragments of the target genes were cloned into the pGreenII 0800-LUC reporter vector, whereas the CDSs of *VvERF5*, *VvERF104*, and *VvMYB14* were inserted into the pGreenII 62-SK effector vector. Through *Agrobacterium*-mediated transformation, specified plasmid combinations were transiently introduced into tobacco leaves. The relative LUC/Renilla (REN) activity was then measured using Dual-LUC assay reagents (Promega, Wisconsin, USA).

### Stable and transient transformations of grape seeds, calli, and/or *Arabidopsis* plants

The open reading frames (ORFs) of various genes were cloned into the pRI101-GUS or pHB vectors to create constructs for overexpression, specifically 35S::VvERF5, 35S::VvERF104, and 35S::VvMYB14. For antisense suppression, the 3′-untranslated regions (3′-UTRs) of *VvERF5*, *VvASMT*, *VvERF104*, and *VvMYB14* were inserted into the pHB vector. The pRI101-GUS vectors containing ORFs were transiently transformed into grape seeds by using the *Agrobacterium tumefaciens* strain LBA4404. In this process, grape seeds were halved lengthwise, soaked in an *Agrobacterium* suspension with gentle shaking for 20 min, and then subjected to vacuum infiltration for 30 min. The seeds were cocultivated on solid MS medium (Murashige and Skoog) containing 2% sucrose and 15 mg·L^−1^ acetosyringone at 25°C in the dark for 3 days.

For the genetic transformation of grape calli, the pHB vectors containing ORFs and 3′-UTRs were introduced for sense overexpression and antisense suppression, respectively, using *Agrobacterium*-mediated transformation (Xu et al., 2019). Calli were immersed in an *Agrobacterium* suspension for 20 min, blotted dry, and transferred to the MS medium containing 100 µM acetosyringone. After 2 days of coculture in darkness at 25°C, the calli were screened on B5 medium containing 250 mg·L^−1^ cefotaxime and 20 mg·L^−1^ hygromycin at 25°C.

To identify the MTRE, pCAMBIA1391 vectors with the 35S promoter replaced by different fragments of *VvMYB14* were transiently transformed into grape calli through *Agrobacterium*-mediated transformation. The calli were soaked in an *Agrobacterium* solution with gentle shaking for 20 min, followed by vacuum infiltration for 10 min. Cocultivation was performed on solid MS medium containing 50 µM melatonin and 15 mg·L^−1^ acetosyringone for 2 days in the dark.

For *Arabidopsis* transformation, the pHB vectors containing ORFs were introduced into *Arabidopsis thaliana* Columbia-0 (Col-0) via the *Agrobacterium* strain GV3101 using the floral dip method (Clough and Bent, 2008).

All transgenic seeds, calli, and *Arabidopsis* plants were confirmed using β-glucuronidase (GUS) staining, polymerase chain reaction (PCR), and/or reverse transcription quantitative PCR (RT-qPCR). All primers were listed in Table S1

### GUS staining and activity assays

Protein content was measured using a bicinchoninic acid (BCA) protein assay kit (Beyotime, Shanghai, China). GUS histochemical staining and activity were assessed using a GUS staining kit, following the manufacturer’s instructions (Coolaber Science & Technology Co., Ltd., Beijing, China).

### RNA-Seq analyses

Total RNA was extracted using TRIzol reagent (Invitrogen, Carlsbad, CA, USA). mRNAs were then purified using poly-T oligo-attached magnetic beads. Sequencing libraries were constructed using the NEBNext Ultra RNA Library Prep Kit for Illumina (#7530 L, NEB, USA) following the manufacturer’s instructions. After preparation, the libraries were sequenced on the Illumina HiSeq 4000 platform, producing 150-bp reads. Grape genome data and corresponding annotations were downloaded from the CRIBI website (http://genomes.cribi.unipd.it/grape/). Transcriptome assembly and quantification were performed using StringTie software (v. 2.1.3b). The expression levels of unigenes were quantified in terms of fragments per kilobase of transcript per million mapped reads (FPKM). The differentially expressed genes (DEGs) were screened using the follow criteria: log2(fold change) ≥1 and false discovery rate (FDR) ≤ 0.05. Data analysis, including Principal component analysis (PCA), volcano plots and Kyoto Encyclopedia of Genes and Genomes (KEGG) enrichment analysis, was conducted using R software (v. 3.5.0) and BLASTALL, respectively.

### Targeted metabolomics analysis of phenolic compounds

Metabolite extraction, analysis, and determination were performed by OE Biotech Co., Ltd. (Shanghai, China). Briefly, lyophilized grape seeds were ground using a mixer mill (MM 400, Retsch). Metabolites were then extracted using 400 µL of chloroform and 600 µL of 70% methanol, followed by centrifugation and filtration. The resulting filtrate was analyzed using an ultra-performance liquid chromatography–electrospray ionization–tandem mass spectrometry (UPLC–ESI–MS/MS) system, which included an ExionLC UPLC and a QTRAP 6500+ mass spectrometer (AB SCIEX, Darmstadt, Germany). The UPLC and MS conditions were set according to the methods of our previous study (Gao et al., 2022). The quantification of phenolic compounds was performed using an external calibration curve made with different phenolic compound standards. Differentially accumulated metabolites (DAMs) were considered based on fold change (FC) ≥ 2 and P-value ≤ 0.005.

### DAP-Seq analysis

For DNA affinity purification sequencing (DAP-seq), genomic DNA from grapes was extracted, purified, and fragmented into approximately 200-bp segments to construct a gDNA library. The CDS of *VvMYB14* was cloned into the pFN19K HaloTag vector. Purification of VvMYB14 and enrichment of its DNA targets were conducted by Bluescape Biotechnology Co. (Baoding, China). Clean reads were mapped to the grape genome by using Bowtie2 software (Langmead and Salzberg, 2012). *VvMYB14* binding peaks were identified using model-based analysis for ChIP-Seq (MACS2) software, with a peak defined by a q value of <0.05 (Yu et al., 2015).

### Real-time quantitative PCR

Total RNA was extracted using an RNAprep Pure Plant Kit (Tiangen, Beijing, China). The extracted RNA was then reverse transcribed into complementary DNA (cDNA) by using the Hiscript Q RT SuperMix (Vazyme, Nanjing, China). Quantitative PCR (qPCR) was performed using ChamQ SYBR qPCR Master Mix (Vazyme, Nanjing, China) on an ABI7500 quantitative real-time PCR (qRT-PCR) instrument (ABI, MA, USA). The primers used for the qRT-PCR are listed in Table S1.

### Statistical analysis

Statistical analysis was performed using SPSS (V19.0) Statistics software. The significance of differences was determined by one-way ANOVA (P<0.05) followed by Tukey’s test. The student’s t-test with a two-tailed distribution was used to compare two sample groups.

## Results

### Ethylene increases melatonin synthesis via VvERF5-induced *VvASMT* expression

To explore the relationship between ethylene and melatonin during grape seed ripening, changes in ethylene production and melatonin content were monitored. Ethylene release peaked at 70 DAB and remained low afterward, except for a minor peak at 100 DAB, continuing until ripening. By contrast, melatonin levels began to rise steadily from 88 DAB, reaching their maximum in ripened seeds (Fig. 1A). This finding indicates that the peak in ethylene production occurred earlier than the increase in melatonin content. Furthermore, exogenous treatment with ethephon increased both the ethylene release rate and melatonin accumulation, whereas treatment with 1-MCP reduced the melatonin content in seeds from 3 to 12 days after treatment (DAT; Fig. 1B, C). Ethephon treatment also upregulated the expression of the melatonin synthesis-related gene *VvASMT*, whereas 1-MCP led to the opposite effect. However, neither ethephon treatment nor 1-MCP treatment led to significant changes in the expression of other melatonin synthesis-related genes, *VvSNAT* and *VvCOMT* (Fig. 1D-F). These findings suggest that ethylene promotes melatonin synthesis by increasing *VvASMT* expression.

**Figure. 1.**
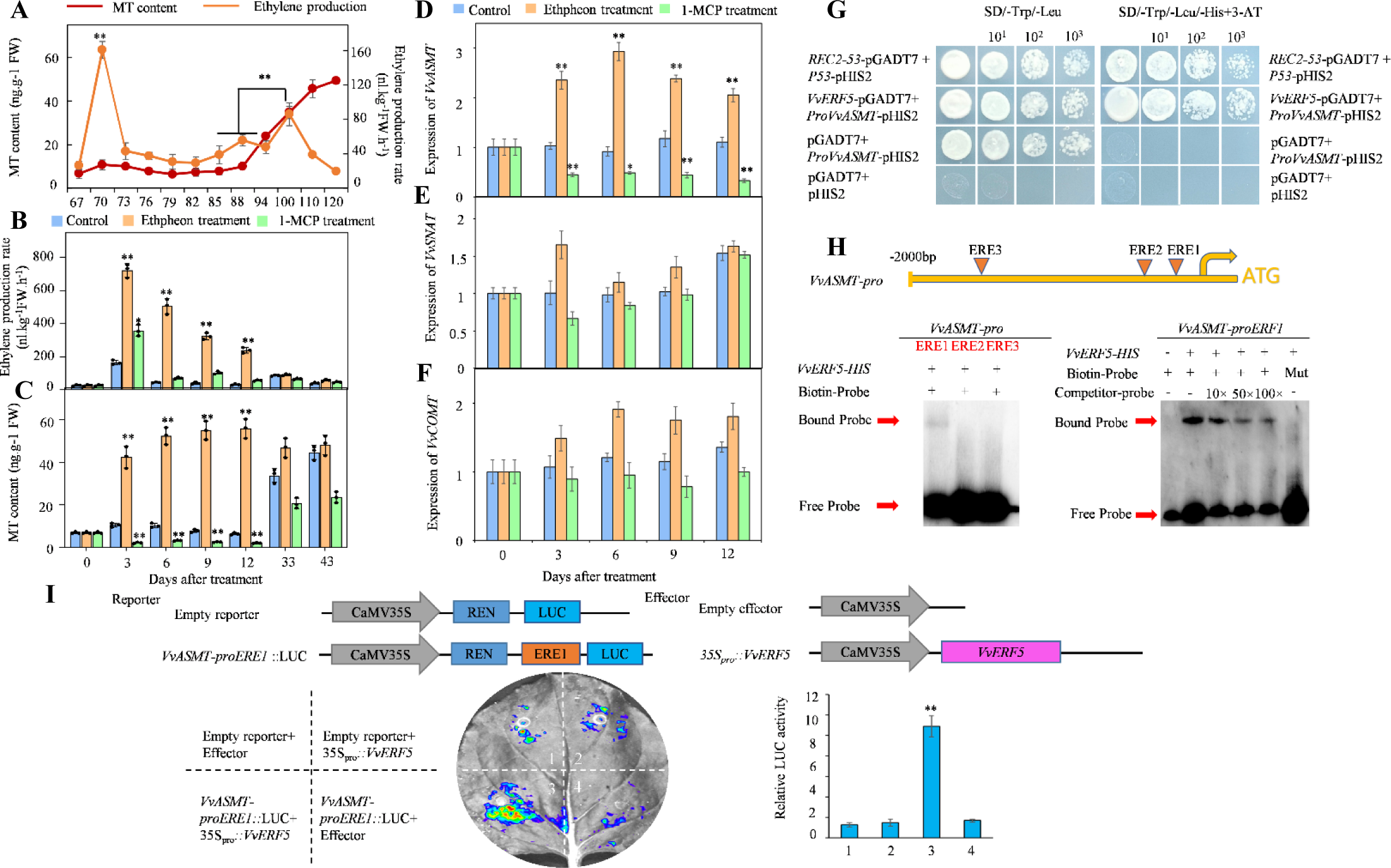
Screening of ethylene-induced melatonin synthesis-related genes and their upstream transcription factors. **A)** Changes in the melatonin content and ethylene production rate during grape seed ripening. **B to F)** Effects of ethephon and 1-MCP treatments on the ethylene production rate, melatonin content, and the expression of *VvASMT*, *VvSNAT*, and *VvCOMT* in grape seeds. **G)** Yeast one-hybrid (Y1H) assay showing the binding of *VvERF5* to ERE1 in the *VvASMT* promoter. **H)** Electrophoretic mobility shift assays demonstrating the binding of *VvERF5* to ERE1, ERE2, and ERE3 within the *VvASMT* promoter. **I)** Fluorescence observations from the dual-luciferase (LUC) assay and relative LUC activity measurements. The data are presented as the means ± SDs of three replicates in panels a-f. ∗, significant difference, P < 0.05; ∗∗, highly significant difference, P < 0.01; MT, melatonin. The sequences of ERE1, ERE2, ERE3 and mERE1 were listed in Table S1.

To identify upstream transcription factors involved in this regulation, a Y1H screening was conducted using ERE1 in the *VvASMT* promoter. This screening identified VvERF5, VvERF016, and VvERF017 as candidate transcription factors. Further analysis revealed that ethephon treatment substantially increased the expression of corresponding genes in grape seeds, whereas 1-MCP treatment led to a decrease (Fig. S1). Subsequent experiments, including Y1H, EMSA, and dual-luciferase (LUC) assays, demonstrated that *VvERF5* specifically bound to ERE1 in the *VvASMT* promoter (Fig. 1G-I) but did not bind to ERE2 or ERE3 (Fig. 1H). In the LUC assays, tobacco leaves cotransformed with the ERE1-35S mini-LUC reporter and the 35S-*VvERF5* effector showed a marked increase in the fluorescence intensity and relative LUC activity, confirming the positive role of *VvERF5* in regulating *VvASMT* expression (Fig. 1I).

To further investigate the role of *VvERF5* in promoting melatonin synthesis, grape seeds were engineered to overexpress *VvERF5* or exhibit suppressed *VvERF5* expression (Fig. 2A, B). Overexpression of *VvERF5* significantly increased both *VvASMT* expression and melatonin content in seeds, whereas suppression of *VvERF5* expression led to the opposite effects (Fig. 2B, C). In addition, three lines of *VvERF5*-overexpressing grape calli (OEC5-1, −2, and −3) and two lines with suppressed *VvERF5* expression (SEC5-1 and −2) were generated (Fig. 2D, E). Consistent with the results in seeds, *VvASMT* expression and melatonin content were significantly increased in calli with gene overexpression and decreased in those with suppressed expression (Fig. 2E-G). The role of *VvERF5* in enhancing *AtASMT* expression and melatonin content was also validated in three lines of *VvERF5*-overexpressing *Arabidopsis* plants (Fig. 2H-J).

**Figure. 2.**
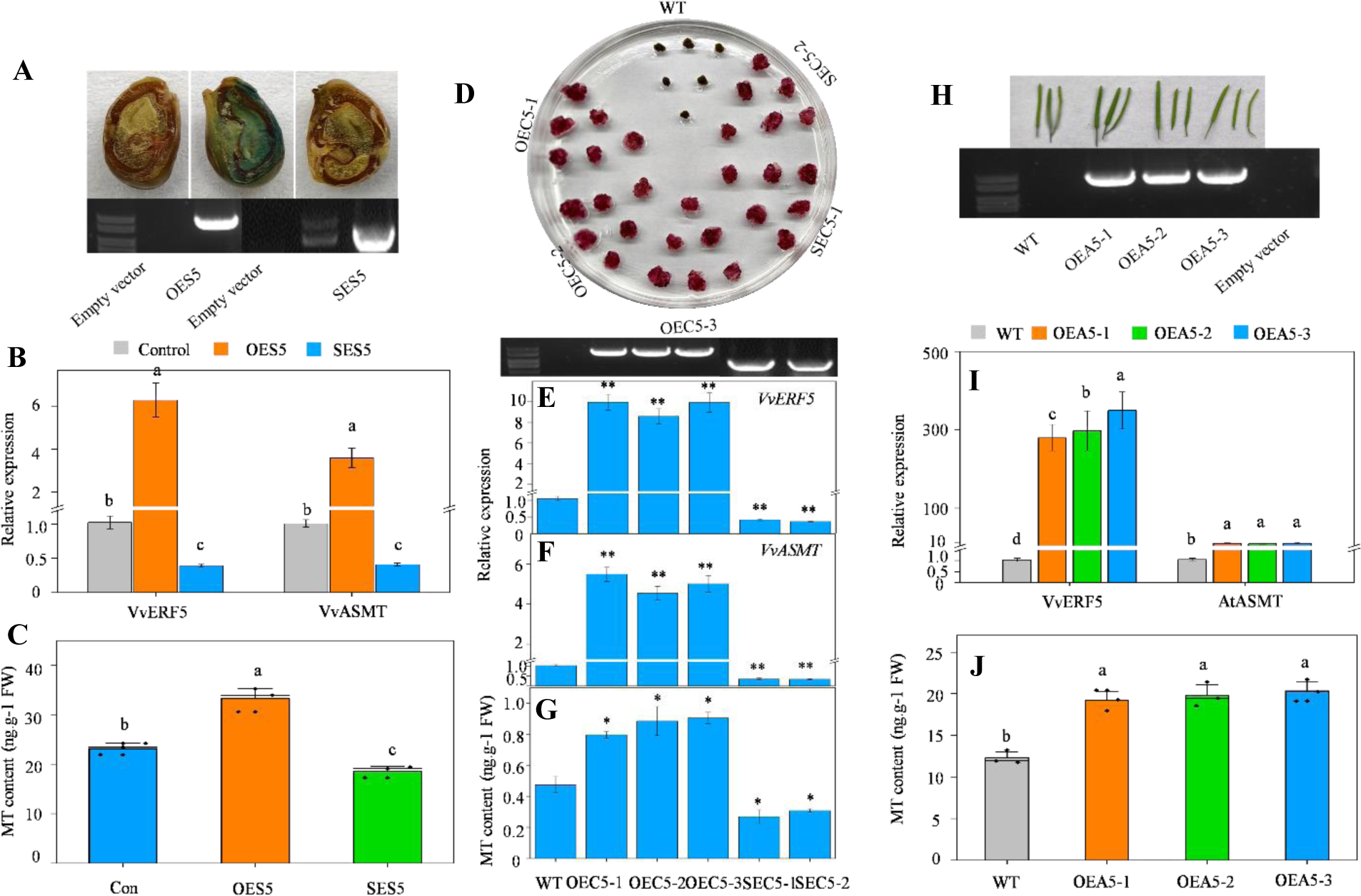
Characterization of the role of *VvERF5* in regulating melatonin synthesis through *VvASMT* expression. **A, B)** Identification of seeds with transient overexpression (OES5) and suppression (SES5) of *VvERF5* using PCR **A)** and qPCR **B)**, and corresponding changes in *VvASMT* expression. Panel (a) shows the construct 35S::ERF5-GUS used to generate seeds overexpressing *VvERF5-GUS*, with GUS staining used for validating the method’s feasibility. **C)** Melatonin content in the control, OES5, and SES5 seeds. **D)** Screening of grape calli overexpressing *VvERF5* and those with suppressed expression using a selective medium. The photograph was taken 25 days after subculture in the selective medium. **E)** Identification of *VvERF5*-overexpressing and -suppressing calli using PCR and qPCR. **F, G)** Expression levels of *VvASMT* **F)** and melatonin content **G)** in WT and transgenic calli. **H)** PCR identification of *VvERF5*-overexpressing *Arabidopsis* plants. **I, J)** Expression levels of *VvERF5* and *AtASMT* **I)**, and melatonin accumulation in *Arabidopsis* seeds. The values represent the means ± SD of three replicates. ∗, Significant difference, P < 0.05; ∗∗, highly significant difference, P < 0.01. The values indicated by the different lowercase letters are significant at P < 0.05; MT, melatonin. The primers used were listed in Table S1.

In summary, ethylene induced the expression of *VvERF5*, which, in turn, transactivated *VvASMT*, leading to increased melatonin biosynthesis.

### Identification of the melatonin-responsive element in the *VvMYB14* promoter

Our previous study demonstrated that the gene *VvMYB14* is strongly induced by melatonin in grape seeds (Zhang et al., 2023). In this study, we selected a 2,268 bp region upstream of the ATG start codon as the *VvMYB14* promoter to identify potential melatonin-responsive elements (MTREs). The promoter was divided into 40 fragments, labeled Pro-1 through Pro-40, which were then inserted into an expression vector to create constructs of each Pro fragment fused to a 35S mini-GUS reporter gene (Fig. 3A). These constructs were transformed into grape calli to evaluate their response to melatonin through GUS staining and GUS activity assays (Fig. 3B). Initially, Pro-1 was found to be induced by melatonin, as indicated by its more intense blue color and higher GUS activity compared with Pro-2. Subsequently, Pro-1 was further divided into two smaller fragments, Pro-3 and Pro-4, with Pro-3 showing strong induction by melatonin. Following this method, a 33-bp fragment, designated Pro-14, was identified as highly responsive to melatonin. Pro-14 was then further divided into 25 shorter fragments, each 8 bp in length. Among these, Pro-24 exhibited the strongest GUS staining intensity and highest activity. By contrast, mutations in one or two base pairs within Pro-24 significantly reduced GUS staining and activity (Fig. 3). Collectively, these results identified Pro-24 (sequence: TGAATATT) as a key MTRE within the *VvMYB14* promoter.

**Figure. 3.**
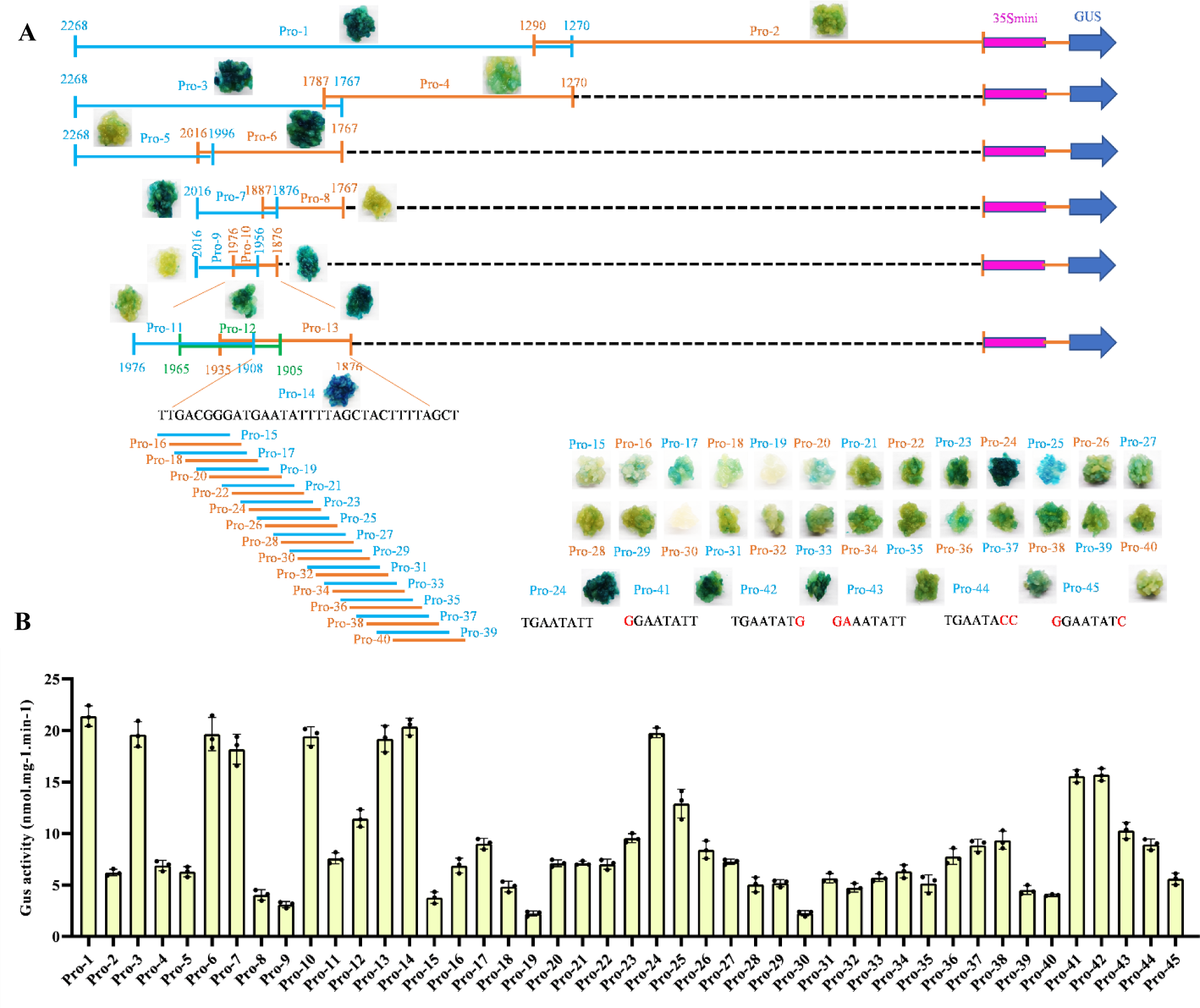
GUS staining **A)** and activity assays **B)** of grape calli expressing the *VvMYB14* promoter fragment-35S mini-GUS constructs. In panel **A)**, different *VvMYB14* promoter fragments are represented by different colored solid lines, with the number indicating the fragment length. Red lines indicate mutant bases in the sequences of Pro-41 to Pro-45. Panel **B)** depicts GUS activity corresponding to these promoter fragments.

### Melatonin-induced *VvERF104* binds to the MTRE in the *VvMYB14* promoter and increases its expression

The *VvMYB14* promoter fragment containing the MTRE was used as bait in a Y1H screening to identify interacting transcription factors. Four transcription factors—VvMYB113, VvWER, VvERF104, and VvERF11—were identified as potential candidates. Among these, all except *VvWER* were significantly induced by melatonin in grape seeds (Fig. S2). Y1H assays further demonstrated that *VvERF104* and *VvERF11* directly bound to the MTRE within the *VvMYB14* promoter (Fig. 4A). The binding of *VvERF104* to the MTRE was confirmed through EMSA and LUC assays. In these assays, *VvERF104* significantly increased the expression of the LUC reporter gene, as indicated by higher luminescence intensity and relative LUC activity. By contrast, *VvERF11* showed weak binding to the MTRE, as evidenced by a faint band in the EMSA and comparable luminescence intensity and relative LUC activity to controls (Fig. 4B, C). Furthermore, overexpression of *VvERF104* in grape seeds led to a significant increase in *VvMYB14* expression, whereas suppression of *VvERF104* resulted in decreased *VvMYB14* expression (Fig. 4D). Similar results were observed in *VvERF104*-overexpression and -suppression grape calli (Fig. 4E). Moreover, overexpression of *VvERF104* in three transgenic *Arabidopsis* lines significantly enhanced the expression of *AtMYB14* in *Arabidopsis* seeds (Fig. 4F). In summary, melatonin induced the expression of *VvERF104*, which in turn transactivated *VvMYB14* by binding to the MTRE in its promoter.

**Figure. 4.**
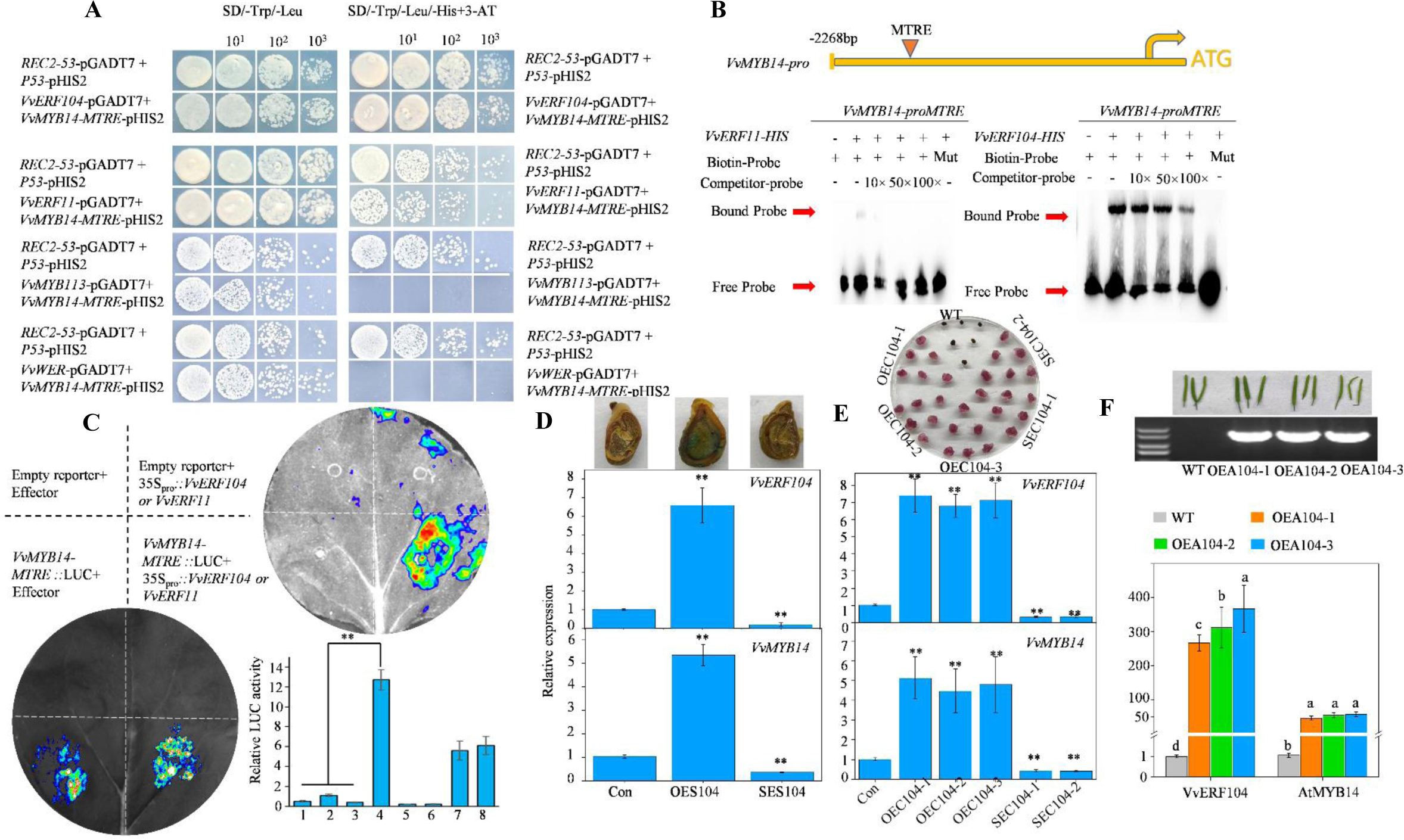
Characterization of the role of *VvERF104* in increasing *VvMYB14* expression by binding to the MTRE. **A)** Yeast one-hybrid (Y1H) assay demonstrating the binding of *VvERF104* and *VvERF11* proteins to the *VvMYB14* promoter. **B)** Electrophoretic mobility shift assays showing the binding of *VvERF104* and *VvERF11* proteins to the MTREs. **C)** Representative images of tobacco leaves 60 h after infiltration, showing corresponding LUC activity. The upper and lower images correspond to the LUC assays of *VvERF104* and *VvERF11*, respectively. **D)** Identification of seeds with transient overexpression (OES104) and suppression (SES104) of *VvERF104* using qPCR and/or GUS staining, along with changes in *VvMYB14* expression in the transgenic seeds. **E)** Identification of *VvERF104*-overexpressing and -suppressing calli using 15 mg·L^−1^ hygromycin selection medium and qPCR, along with changes in *VvMYB14* expression in the transgenic calli. **F)** Identification of *VvERF104*-overexpressing *Arabidopsis* plants using PCR and qPCR, along with changes in *AtMYB14* expression in the transgenic plants. The values represent the means ± SD of three replicates. ∗, Significant difference, P < 0.05; ∗∗, highly significant difference, P < 0.01.The sequences of MTRE and mMTRE were listed in Table S1.

### *VvMYB14* overexpression widely regulates gene expression and metabolite accumulation in the phenylpropanoid pathway

To investigate the function of *VvMYB14*, two groups of *VvMYB14*-overexpressing grape seeds (OES14-1 and OES14-2) were generated (Fig. 5A). RNA-Seq was performed on wild-type (WT), OES14-1, and OES14-2 seeds to evaluate the gene expression changes resulting from *VvMYB14* overexpression. Principal component analysis (PCA) of the nine samples revealed that the first principal component (PC1) accounted for 98.13% of the variance, indicating a substantial difference between WT and *VvMYB14*-overexpressing seeds (Fig. 5B). A total of 3,245 and 3,168 differentially expressed genes (DEGs) were identified in the comparisons of WT vs OES14-1 and WT vs OES14-2, respectively (Fig. 5C, D; Table. S2, S3). KEGG enrichment analysis showed that the differentially expressed genes (DEGs) were primarily associated with flavonoid biosynthesis, followed by phenylpropanoid biosynthesis, phenylalanine metabolism, and plant hormone signal transduction pathways (Fig. 5E, F).

**Figure. 5.**
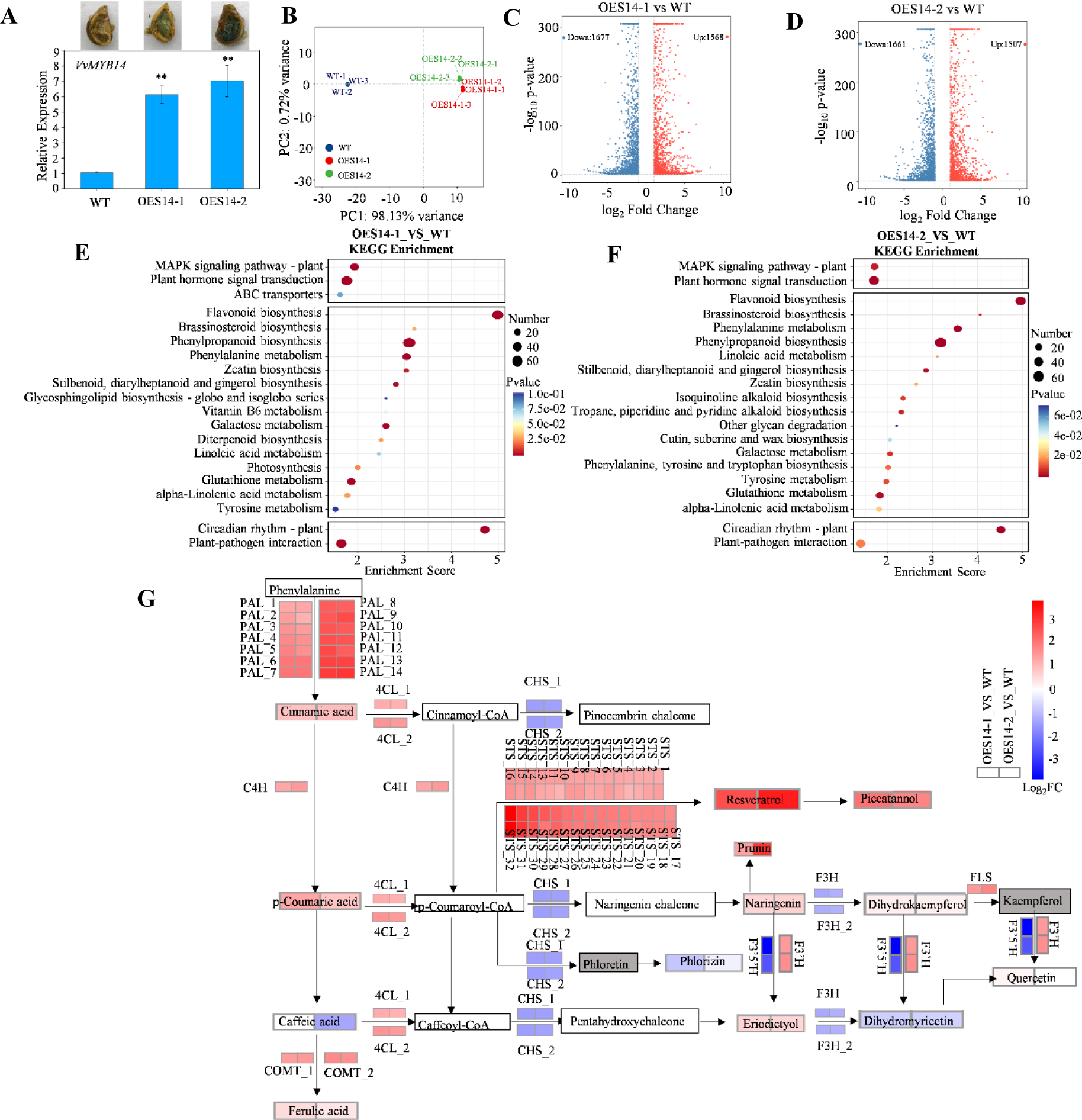
Changes in gene expression and metabolite content in the phenylpropanoid pathway caused by *VvMYB14* overexpression in grape seeds. **A)** Identification of transient overexpression of *VvMYB14* in grape seeds using GUS staining and qPCR. **B)** Principal component analysis (PCA) of the nine samples based on fragments per kilobase of transcript per million mapped reads (FPKM) values. **C, D)** Volcano plots showing upregulated and downregulated genes in the comparison groups OES14-1 vs WT **C)** and OES14-2 vs WT **D)**. **E, F)** KEGG enrichment analysis of differentially expressed genes (DEGs) in OES14-1 **E)** and OES14-2 **F)** compared to WT. **G)** KEGG pathway analysis of metabolites and genes whose accumulation and expression, respectively, were altered in the OE14 seeds compared with the WT seeds. Each solid black arrow represents an enzyme-catalyzed process. White background boxes indicate metabolites not detected in this study, gray boxes represent undetectable metabolites, and colored boxes represent metabolites and DEGs with altered accumulation and expression due to *VvMYB14* overexpression.

Targeted metabolomics was conducted to identify differentially accumulated compounds (DACs), which revealed a total of 130 phenolic compounds; among these, 70 compounds, including 52 phenolic compounds, were present in grape seeds (Table S4). Compared with WT seeds, 17 phenolic compounds were found to be more abundant, whereas 14 were less abundant in the *VvMYB14*-overexpressing seeds (Table S4). An association analysis of DACs and DEGs revealed that the significant upregulation of 14 phenylalanine ammonia-lyases (PALs), one cinnamate-4-hydroxylase (C4H), and two caffeic acid o-methyltransferases (COMTs) dominated the phenylpropanoid biosynthesis pathway, leading to increased levels of cinnamic acid, p-coumaric acid, and ferulic acid (Fig. 5G). Notably, the expression of 32 stilbene synthase (STS) genes significantly increased, which corresponded with elevated levels of resveratrol and piceatannol, leading to increased stilbene biosynthesis.

### VvMYB14 directly binds to the promoters of *VvPAL, VvC4H* and *VvCHS* and regulates their expression

DNA affinity purification sequencing (DAP-seq) was conducted to identify *VvMYB14* binding sites across the grape genome. A total of 10,939 peaks, which were included into 8,769 genes, were uncovered (Table. S5). The binding peaks were predominantly located near the transcription start sites (TSS) of target genes (Fig. 6A), with 18.3% of the peaks found within promoter regions (up to 2.0 kb upstream of the TSS) (Fig. 6B). A total of 1,998 high-confidence VvMYB14 binding sites were discovered, which were located in the promoter regions of 1,739 putative target genes (Table S5). Based on the 29,836 motifs in the promoter regions of 1,935 unique genes (Table. S6), the significantly enriched motifs (E-value < 0.05) were identified using MEME-chip. It was indicated that the MEME-1 motif (DDDDGGTWGGTGRRD) had the highest enrich score, followed by MEME-3 (TTYTYTCTYCTYTCTCTTCTYTCTY), MEME-5 (TTATACCTATAACCAATTATT) and MEME-6 (GRGRGKGGCWTSTCYCCATKRSGWG) (Fig. 6C). KEGG analysis revealed that the putative target genes of *VvMYB14* were highly enriched in pathways related to phenylpropanoid biosynthesis; flavonoid biosynthesis; and stilbenoid, diarylheptanoid, and gingerol biosynthesis (Fig. 6D).

**Figure. 6.**
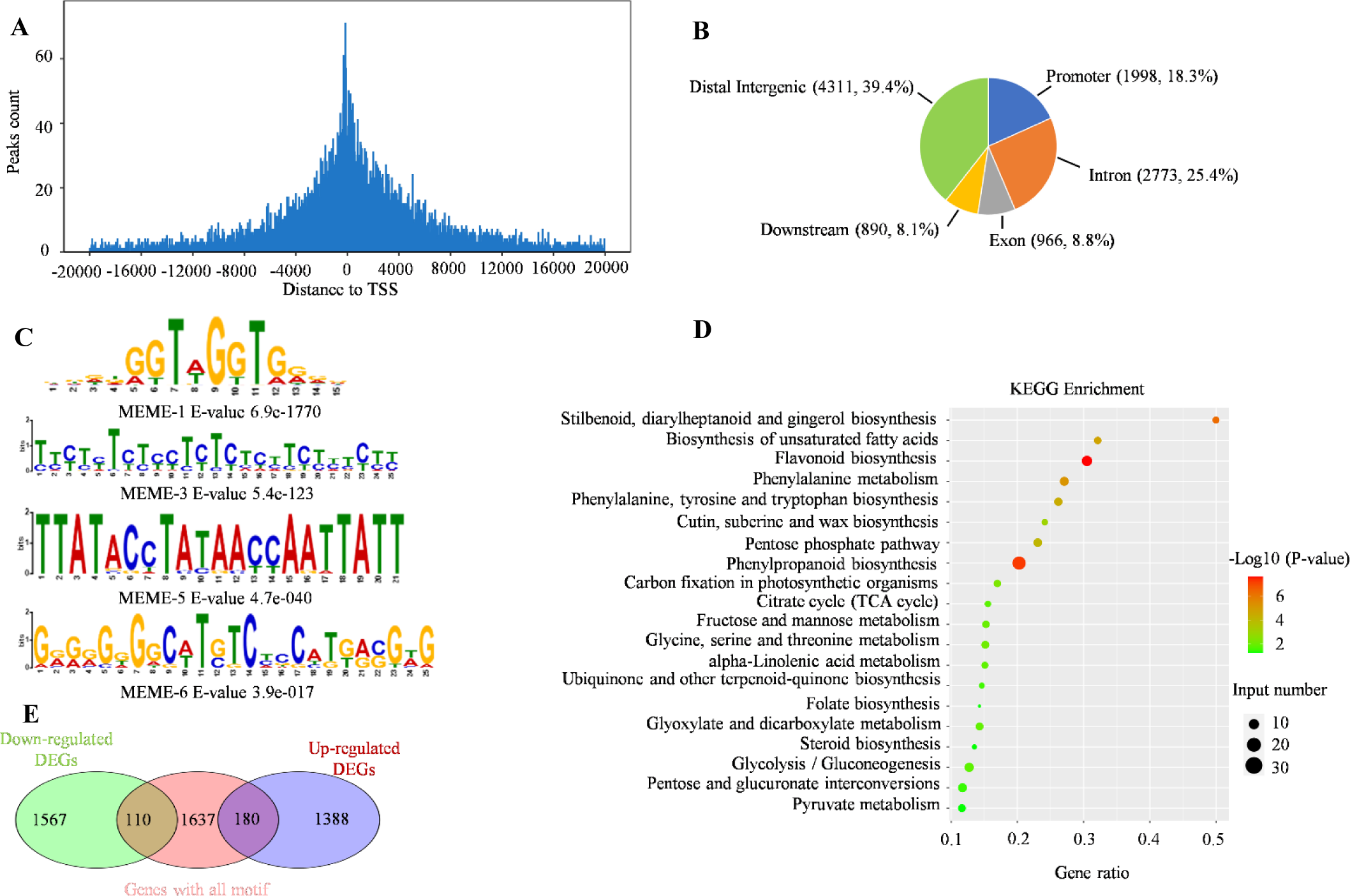
Genome-wide identification of *VvMYB14* binding sites using DNA affinity purification sequencing (DAP-seq). **A)** Distribution of *VvMYB14* binding sites within the −20,000 to +20,000 bp region flanking the transcription start site (TSS). **B)** Number and percentage of peaks in different genomic regions. **C)** Predicted *VvMYB14* binding motif with a high enrichment score. **D)** KEGG analysis of putative target genes of *VvMYB14*. **E)** Venn diagram showing the overlapping genes identified through DAP-seq and RNA-seq.

A combined analysis of DAP-seq and RNA-seq data revealed that 180 upregulated and 110 downregulated DEGs contained *VvMYB14*-binding motifs (Fig. 6E; Table S7). Among these, 78 DEGs were related to phenylpropanoid and flavonoid biosynthesis (Table S7). From this group, 10 DEGs involved in the phenylpropanoid pathway and containing the MEME-1 motif were selected for Y1H assays (Fig. 7A; Fig. S3). Y1H assays demonstrated that *VvMYB14* directly bound to P1 sites in the promoters of *VvPAL* and *VvC4H*, as well as to the P5 site in the promoter of *VvCHS*, with all these sites containing the MEME-1 motif (Fig. 7A, B; Table S1). Further validation using EMSA and LUC analysis confirmed that *VvMYB14* bound to the promoters of *VvPAL*, *VvC4H*, and *VvCHS* (Fig. 7C, D). LUC analysis also revealed that *VvMYB14* increased LUC expression by binding to the P1 site of the *VvPAL* or *VvC4H* promoter and reduced LUC expression by binding to the P5 site of the *VvCHS* promoter (Fig. 7E). qRT-PCR showed that the overexpression of *VvMYB14* led to increased expression of *VvPAL* and *VvC4H* and decreased expression of *VvCHS* in grape seeds, whereas suppression of *VvMYB14* produced the opposite effects (Fig. 7F). Similar results were observed in *VvMYB14*-overexpressing grape calli and those with suppressed expression (Fig. 7G). In summary, these findings indicated that *VvMYB14* regulated the transcription of *VvPAL*, *VvC4H*, and *VvCHS* by binding to the MEME-1 motif in their promoters.

**Figure. 7.**
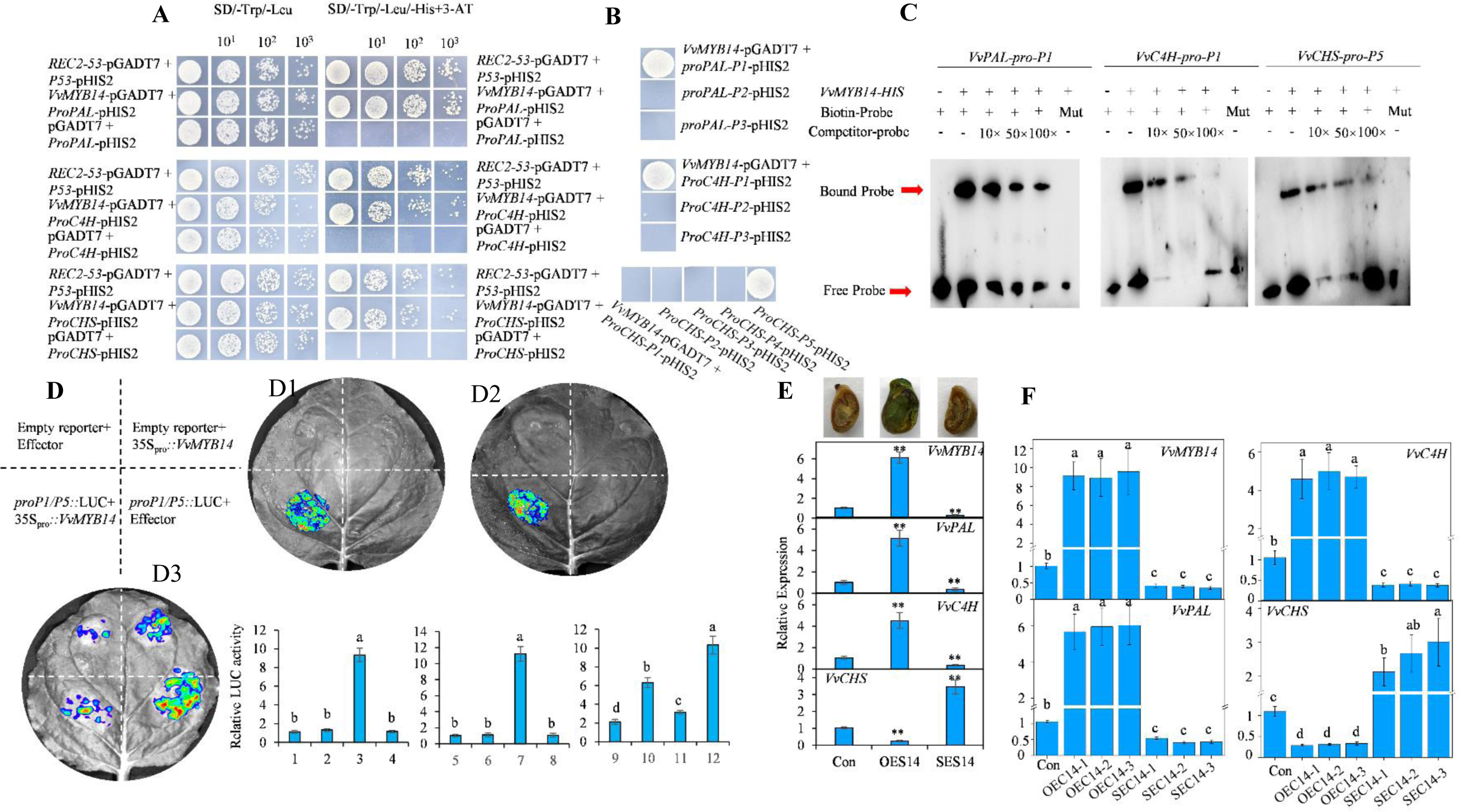
Identification of *VvMYB14* target genes in the phenylpropanoid pathway. **A)** Yeast one-hybrid (Y1H) assays showing the binding of *VvMYB14* to the promoters of *VvPAL*, *VvC4H*, and *VvCHS*, all containing the putative binding sites. The yeast cells diluted 1-, 10-, 100-, and 1000-fold are shown from left to right. **B)** Y1H assays showing the binding of *VvMYB14* to specific sites in the promoters of *VvPAL*, *VvC4H*, and *VvCHS*. **C)** Electrophoretic mobility shift assays (EMSA) showing the binding of *VvMYB14* to cis-regulatory elements in the *VvPAL*, *VvC4H*, and *VvCHS* promoters. **D)** Dual-luciferase assay and LUC activity in tobacco leaves 60 h after infiltration. D1, D2, and D3 indicate the binding of *VvMYB14* to the P1/P5 sites in the promoters of *VvPAL*, *VvC4H*, and *VvCHS*, respectively. **E, F)** Identification of *VvMYB14*-overexpressing and -suppressing seeds **E)** and calli **F)** using GUS staining and/or qRT-PCR, and the effects of altered *VvMYB14* expression on *VvPAL*, *VvC4H*, and *VvCHS* expressions. The values represent the means ± SD of three replicates. ∗, Significant difference, P < 0.05; ∗∗, highly significant difference, P < 0.01.

### ERF5-melatonin-ERF104 pathway participates in ethylene-induced expression of the genes involved in phenylpropanoid pathway

In WT grape seeds, treatments with ethylene and melatonin significantly altered the expression of several key genes, including *VvERF5*, *VvASMT*, *VvERF104*, *VvMYB14*, *VvPAL*, *VvC4H*, and *VvCHS* (Fig. S4). By contrast, suppression of *VvERF5*, *VvASMT*, or *VvERF104* mitigated the ethylene-induced increases in the expression of *VvMYB14*, *VvPAL*, and *VvC4H* while enhancing the ethylene-induced expression of *VvCHS*. These results underscored the critical role of these genes in ethylene signaling. Moreover, the suppression of *VvASMT* reduced the ethylene-induced expression of these genes, indicating that ethylene’s effects are partially mediated through melatonin. Based on these findings, the regulatory pathway of ERF5-melatonin-ERF104 was proposed to participate in the regulation of ethylene on phenylpropanoid pathway.

## Discussion

### VvERF5 might regulate the phenylpropanoid pathway via multiple pathways including melatonin signaling

Grapes, as a typical nonclimacteric fruit, do not typically exhibit a peak in ethylene release during ripening. However, ethylene release peaks have been detected in grape varieties, such as Cabernet Sauvignon, Moldova, and Muscat Hamburg (Chervin et al., 2004; Sun et al., 2010; Xu et al., 2018). In this study, an ethylene release peak was observed in Merlot grape seeds at 70 DAB (Fig. 1A). This suggests that significant ethylene production occurs in various grape tissues before the onset of ripening, which precedes the peak in melatonin accumulation during berry ripening (Fig. 1a; 22). Furthermore, ethylene treatment was found to increase melatonin synthesis, whereas treatment with 1-MCP produced the opposite effect (Fig. 1C). These findings suggest that ethylene may trigger or at least modulate melatonin synthesis during fruit development.

ERFs are downstream components of the ethylene signaling pathway that regulate the expression of ethylene-responsive genes by directly binding to their promoter regions (Pirrello et al., 2012). In this study, *VvERF5* expression in the seeds increased after ethephon treatment but decreased with 1-MCP treatment (Fig. S1). Similar to the role of *SlERF5* in tomatoes, which induces the ethylene-responsive “triple response” phenotype (Pan et al., 2011), *VvERF5* appears to play a crucial role in the ethylene signaling pathway. Our results indicated that *VvASMT* is a target gene of *VvERF5* (Fig. 1G-I). *ASMT* has been shown to play a rate-limiting role in melatonin synthesis in capsicum (Pan et al., 2019; Wu et al., 2022). This suggests that *VvERF5* regulates melatonin synthesis via *VvASMT*. This hypothesis was further supported by the effects of *VvERF5* overexpression and suppression on *VvASMT* expression and melatonin content in the seeds and calli (Fig. 2A-G). Melatonin’s role in altering the metabolism of secondary metabolites, primarily derived from the phenylpropanoid pathway, has been reported (Jayarajan and Sharma, 2021; Fan et al., 2022). Aligning with these findings, our study demonstrated that the suppression of *VvERF5* reduced ethylene-induced expression of *VvPAL* and *VvC4H* (Fig. S4), indicating that *VvERF5* regulates the phenylpropanoid pathway via melatonin signaling.

*ERF5* has been reported to regulate flavonoid biosynthesis through other pathways. In pear, *PcERF5* activates the anthocyanin synthesis-related transcription factors *PcMYB10* and *PcMYB114*, as well as *MYBA* in mulberry (Mo et al., 2022; Chang et al., 2023). It also transactivates flavonoid synthesis-related genes, including *DFR*, *ANS*, *UFGT*, and *F3H* (Mo et al., 2022; Chang et al., 2023). In addition, *PcERF5* interacts with *PcMYB10* to form the ERF5–MYB10 protein complex, which enhances the transcriptional activation of *PcERF5* on its target genes (Chang et al., 2023). In summary, *VvERF5* is a key component of the ethylene signaling cascade and plays a role in regulating the phenylpropanoid pathway via melatonin signaling and other mechanisms. Increasing evidence suggests that melatonin promotes ethylene biosynthesis by upregulating the expression of *ACS* and/or *ACO* in fruits, including grapes (Xu et al., 2018; Sun et al., 2020; Ma et al., 2021). This finding indicates the potential existence of a regulatory circuit between ethylene and melatonin synthesis, which may contribute to the fine-tuning of the phenylpropanoid pathway.

### *VvMYB14* broadly regulates the phenylpropanoid pathway possibly by binding to different target genes

Overexpression of *VvMYB14* in grape seeds resulted in significant changes in the expression of genes involved in the phenylpropanoid pathway and altered the content of 31 phenolic compounds compared with WT seeds (Fig. 5E, F; Table S4). Specifically, a combined analysis of DAP-seq and RNA-seq revealed the presence of *VvMYB14* binding motifs in the promoter regions of 78 DEGs associated with the phenylpropanoid pathway (Fig. 6E; Table S7). Similarly, overexpression of *LiMYB14* in lotus plants increased the expression of genes involved in the general phenylpropanoid pathway, including *PAL*, *C4H*, and *4CL* (Shelton et al., 2012). These findings suggest that *MYB14* broadly modulates the phenylpropanoid pathway by regulating different target genes. Additionally, *VvPAL*, *VvC4H*, and *VvCHS* were confirmed as direct targets of *VvMYB14* (Fig. 7F, G).

PALs catalyze the first step of the phenylpropanoid pathway, converting phenylalanine into cinnamic acid (Barros and Dixon, 2020). Cinnamic acid serves as the initial substrate for the biosynthesis of other phenylpropanoids and phenolic compounds (Huang et al., 2010). In this study, overexpression of *VvMYB14* led to an increase in the expression of 14 *VvPAL* genes and the content of cinnamic acid (Fig. 5G). Notably, 13 of these *VvPAL* genes also contained the MEME-1 motif in their promoters, suggesting that they might be direct targets of *VvMYB14* (Table S7). This indicates that *VvMYB14* could induce the entire phenylpropanoid pathway by enhancing the production of key initial substrates. Cinnamic acid is hydroxylated by trans-cinnamate 4-hydroxylase (C4H) to produce p-coumaric acid (Barros and Dixon, 2020), and increased expression of *VvC4H* contributed to higher levels of p-coumaric acid in the overexpressing seeds (Fig. 5G). Chalcone synthase (CHS) catalyzes the first committed step in flavonoid biosynthesis by directing carbon flux from general phenylpropanoid metabolism to the flavonoid pathway (Zhang et al., 2017). The downregulation of two *VvCHS* genes suggests that more substrates may be diverted to other branches, such as resveratrol biosynthesis (Fig. 5G). Ethylene treatment and *VvMYB14* overexpression had different effects on *VvCHS* expression (Fig. 7F; Fig. S4), suggesting that additional regulators are involved in controlling *VvCHS* expression within the ethylene signaling pathway.

*VvSTS41* and *VvSTS29* have been identified as target genes of *VvMYB14* (Höll et al., 2013). The significant increase in the expression of 17 *VvSTS* genes, along with higher levels of resveratrol and piceatannol (Fig. 5G), indicates the crucial role of *VvMYB14* in regulating resveratrol synthesis through the activation of *VvSTS*. Moreover, *VvMYBPA1* is directly transactivated by *VvMYB14*, leading to increased proanthocyanidin synthesis in grape seeds (Zhang et al., 2023). In *Medicago truncatula*, *MtMYB14* and *MtMYB5* physically interact and synergistically activate the expression of anthocyanidin reductase (ANR) and leucoanthocyanidin reductase (LAR) (Liu et al., 2014). *VvMYB14* lacks the motif necessary for interaction with basic helix-loop-helix (bHLH) proteins, suggesting that *VvMYB14* induces promoter activity of target genes independently of bHLH/WD40 cofactors (Höll et al., 2013; Zhang et al., 2023), suggesting that *VvMYB14* regulates the phenylpropanoid pathway by directly controlling the expression of multiple target genes. Nevertheless, the expression of *HST, 4CL2, CCOAOMT, CCR1, COMT, CYP98A2* and *CAD1*, were largely changes by *VvMYB14* overexpression but they were not its target genes (Table. S2, S3), suggesting that *VvMYB14* regulates gene expression indirectly via other pathways.

### VvERF104 might integrate ethylene and melatonin signals to regulate the phenylpropanoid pathway

Ethylene and melatonin act as regulators of the phenylpropanoid pathway in grapes by modulating gene expression. In this study, both ethylene and melatonin treatments led to an increase in the expression of key genes involved in the phenylpropanoid pathway, including *VvPAL*, *VvC4H*, and *VvCHS* (Fig. S4). Similar findings have been reported in previous studies where ethylene or melatonin treatment upregulated the phenylpropanoid pathway genes, including *VvPAL*, *VvC4H*, and *VvCHS* (Sharafi et al., 2021; Wang et al., 2022). However, the precise mechanisms by which ethylene and melatonin regulate this pathway remain largely unknown. Here, we identified VvERF104 as a key transcription factor that is strongly induced by both ethylene and melatonin. *VvERF104* was shown to bind directly to the MTRE in the promoter of *VvMYB14* (Fig. S2; Fig. 4A-C). Additionally, suppression of *VvERF104* significantly reduced the ethylene- or melatonin-induced expression of *VvMYB14*, *VvPAL*, and *VvC4H* (Fig. S4). These findings suggest that *VvERF104* plays a crucial role in integrating ethylene and melatonin signals to regulate the phenylpropanoid pathway.

The role of ERF104 in integrating ethylene and melatonin signals to induce immunity in *Arabidopsis thaliana* has been previously reported. Specifically, the flg22 signaling network induces MPK6 to directly target ERF104 through phosphorylation, affecting ERF104 stability and ethylene signaling. Simultaneously, MPK3/6 and MKK4/5 stimulate ethylene production, which triggers the release of MPK6 from ERF104 in a process dependent on EIN2 and the EIN3/EIL members. The liberated ERF104 then enhances immunity by regulating its target genes, positioning ERF104 as a key regulator of basal immunity in *Arabidopsis (Bethke et al., 2009)*. Melatonin has also been shown to increase the expression and phosphorylation levels of MPK3/6, which, in turn, activate several transcription factors that induce various defense genes in *Panax notoginseng* and *A. thaliana* (Lee and Back, 2016; Yang et al., 2021). These findings suggest that melatonin may induce immunity via the MPK3/6-ERF104 pathway. It is worth investigating whether a similar mechanism exists in the regulation of phenylpropanoid metabolism by ethylene and melatonin.

In this study, *VvERF104* bound to the MTRE in the promoter of *VvMYB14* and increased its expression (Fig. 4A-C). Genome-wide MTRE assays revealed that this element exists in the promoters of 90 other genes related to the phenylpropanoid pathway (Table S8). Additionally, our previous work demonstrated that *VvERF104* directly transactivates the expression of *VvMYBPA1* (Zhang et al., 2023), suggesting that *VvERF104* may regulate the phenylpropanoid pathway by controlling transcription factors or structural genes, other than *VvMYB14*.

In summary, ethylene promotes melatonin biosynthesis by inducing the expression of *VvERF5*, which transactivates *VvASMT*. Melatonin, in turn, strongly induces *VvMYB14*, which broadly modulates gene expression and metabolite content in the phenylpropanoid pathway in grape seeds. *VvMYB14* binds to the MEME-1 motif in the promoters of *VvPAL*, *VvC4H*, and *VvCHS* to regulate their expression, making it a key transcription factor in melatonin’s regulation of the phenylpropanoid pathway. Additionally, the MTRE was identified within the *VvMYB14* promoter, and *VvERF104* was shown to bind to the MTRE and activate *VvMYB14* expression. The results also indicate the roles of *VvERF5*, *VvASMT*, and *VvERF104* in mediating ethylene-induced expression of genes involved in the phenylpropanoid pathway. Based on these results, we propose an ERF5-melatonin-ERF104 regulatory pathway that explains how ethylene regulates the phenylpropanoid pathway via *VvMYB14* (Fig. 8).

**Figure. 8.**
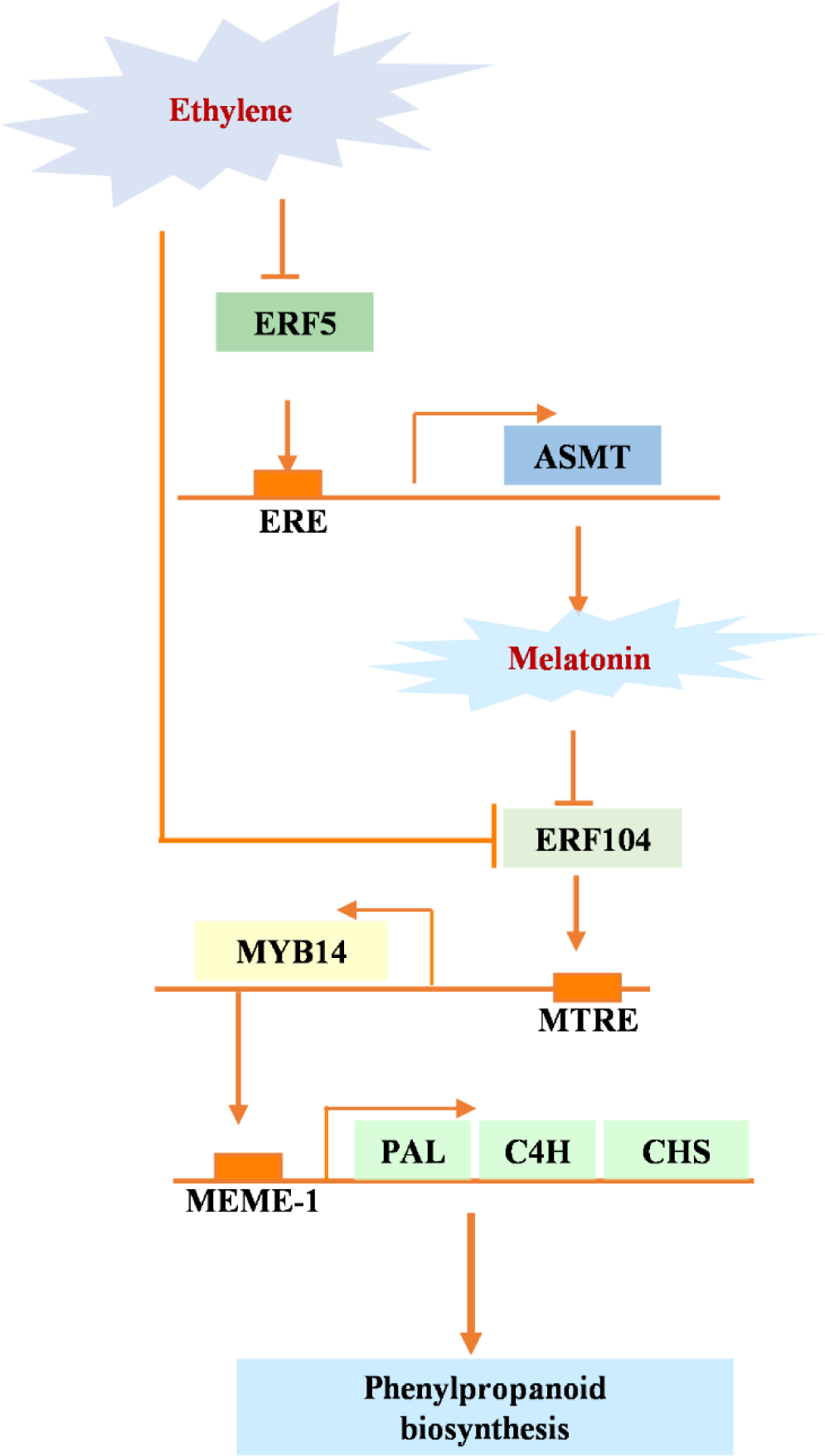
Model of phenylpropanoid pathway regulation by ethylene via the ERF5-Melatonin-ERF104 pathway. In this model, *VvERF5* transactivates *VvASMT* to increase melatonin synthesis. Ethylene promotes *VvERF104* expression directly or indirectly through melatonin. *VvERF104* then binds to the MTRE in the *VvMYB14* promoter to induce its expression. *VvMYB14* regulates the expression of *VvPAL*, *VvC4H*, and *VvCHS* by binding to the MEME-1 motif.

## Acknowledgments

We are highly thankful to Professor Yuanpeng Du and Zhen Gao for their helpful suggestions.

## Author contributions

Y.Y. and G.S. conceived and designed the research; G.S., W.F., W.S., and L.S. performed the experiments; X.Y., K.H. and L.X. analyzed the data; G.S. and Y.Y. wrote the manuscript. All authors read and approved the manuscript.

## Supplemental data

**Supplemental Figure S1** Changes in the expression of screened genes in control grape seeds and those treated with ethephon or 1-MCP at different days after treatment.

**Supplemental Figure S2** Changes in the expression of screened genes in control and melatonin-treated grape seeds at different days after treatment.

**Supplemental Figure S3** Yeast one-hybrid assays of VvMYB14 with seven possible target genes. The yeast cells were diluted 1-, 10-, 100-, and 1000-fold, respectively, shown from left to right; 3-AT was used as a screening marker.

**Supplemental Figure S4** Changes in gene expression in the control and ethephon- or melatonin-treated grape seeds with suppressed expression of *VvERF5*, *VvASMT*, and *VvERF104*.

**Supplemental Table S1** Sequences of elements and primers in this study

**Supplemental Table S2** DEGs between OES14-1 and WT

**Supplemental Table S3** DEGs between OES14-2 and WT

**Supplemental Table S4** All of the metabolites detected in the WT and OES14-1/OES14-2 seeds

**Supplemental Table S5** Binding peaks of VvMYB14 detected using DAP-seq analysis.

**Supplemental Table S6** The determination of the binding motifs in the peaks.

**Supplemental Table S7** Overlapping genes of RNA-seq and DAP-seq

**Supplemental Table S8** All of the genes contain MTRE in promoter

## Data availability

The full RNA-seq and DAP-seq data have been submitted to the Sequence Read Archive (SRA) of the NCBI under BioSample accession PRJNA1151140 and PRJNA1151207.

## References

Arc E, Sechet J, Corbineau F, Rajjou L, Marion-Poll A (2013) ABA crosstalk with ethylene and nitric oxide in seed dormancy and germination. Frontiers in Plant Science 4

Arnao MB, Hernández-Ruiz J (2019) Melatonin: A New Plant Hormone and/or a Plant Master Regulator? Trends in Plant Science 24: 38–48

Back K, Tan DX, Reiter RJ (2016) Melatonin biosynthesis in plants: multiple pathways catalyze tryptophan to melatonin in the cytoplasm or chloroplasts. Journal of Pineal Research 61: 426–437

Bakshi A, Shemansky JM, Chang C, Binder BM (2015) History of Research on the Plant Hormone Ethylene. Journal of Plant Growth Regulation 34: 809–827

Barros J, Dixon RA (2020) Plant Phenylalanine/Tyrosine Ammonia-lyases. Trends in Plant Science 25: 66–79

Becatti E, Genova G, Ranieri A, Tonutti P (2014) Postharvest treatments with ethylene on Vitis vinifera (cv Sangiovese) grapes affect berry metabolism and wine composition. Food Chemistry 159: 257–266

Bethke G, Unthan T, Uhrig JF, Pöschl Y, Gust AA, Scheel D, Lee J (2009) Flg22 regulates the release of an ethylene response factor substrate from MAP kinase 6 in *Arabidopsis thaliana* via ethylene signaling. Proceedings of the National Academy of Sciences of the United States of America 106

Caspi Y, Pantazopoulou CK, Prompers JJ, Pieterse CMJ, Hulshoff Pol H, Kajala K (2023) Why did glutamate, GABA, and melatonin become intercellular signalling molecules in plants? eLife 12

Chang Y-j, Chen G-s, Yang G-y, Sun C-r, Wei W-l, Korban SS, Wu J (2023) The PcERF5 promotes anthocyanin biosynthesis in red-fleshed pear (Pyrus communis) through both activating and interacting with PcMYB transcription factors. Journal of Integrative Agriculture 22: 2687–2704

Chervin C, El-Kereamy A, Roustan J-P, Latché A, Lamon J, Bouzayen M (2004) Ethylene seems required for the berry development and ripening in grape, a non-climacteric fruit. Plant Science 167: 1301–1305

Clough SJ, Bent AF (2008) Floral dip: a simplified method forAgrobacterium-mediated transformation ofArabidopsis thaliana. The Plant Journal 16: 735–743

Fan S, Li Q, Feng S, Lei Q, Abbas F, Yao Y, Chen W, Li X, Zhu X (2022) Melatonin Maintains Fruit Quality and Reduces Anthracnose in Postharvest Papaya via Enhancement of Antioxidants and Inhibition of Pathogen Development. Antioxidants 11

Gao S, Wang F, Zhang X, Li B, Yao Y (2022) Characterization of anthocyanin and nonanthocyanidin phenolic compounds and/or their biosynthesis pathway in red-fleshed ‘Kanghong’ grape berries and their wine. Food Research International 161

Hanlin RL, Kelm MA, Wilkinson KL, Downey MO (2011) Detailed Characterization of Proanthocyanidins in Skin, Seeds, and Wine of Shiraz and Cabernet Sauvignon Wine Grapes (Vitis vinifera). Journal of Agricultural and Food Chemistry 59: 13265–13276

Höll J, Vannozzi A, Czemmel S, D’Onofrio C, Walker AR, Rausch T, Lucchin M, Boss PK, Dry IB, Bogs J (2013) The R2R3-MYB Transcription Factors MYB14 and MYB15 Regulate Stilbene Biosynthesis in Vitis vinifera. The Plant Cell 25: 4135–4149

Huang J, Gu M, Lai Z, Fan B, Shi K, Zhou Y-H, Yu J-Q, Chen Z (2010) Functional Analysis of the Arabidopsis PAL Gene Family in Plant Growth, Development, and Response to Environmental Stress. Plant Physiology 153: 1526–1538

Huang Q, Sun M, Yuan T, Wang Y, Shi M, Lu S, Tang B, Pan J, Wang Y, Kai G (2019) The AP2/ERF transcription factor SmERF1L1 regulates the biosynthesis of tanshinones and phenolic acids in Salvia miltiorrhiza. Food Chemistry 274: 368–375

Jayarajan S, Sharma RR (2021) Melatonin: A blooming biomolecule for postharvest management of perishable fruits and vegetables. Trends in Food Science & Technology 116: 318–328

Jie H, He P, Zhao L, Ma Y, Jie Y (2023) Molecular Mechanisms Regulating Phenylpropanoid Metabolism in Exogenously-Sprayed Ethylene Forage Ramie Based on Transcriptomic and Metabolomic Analyses. Plants 12

Langmead B, Salzberg SL (2012) Fast gapped-read alignment with Bowtie 2. Nature Methods 9: 357–U354

Lee HY, Back K (2016) Mitogen-activated protein kinase pathways are required for melatonin-mediated defense responses in plants. Journal of Pineal Research 60: 327–335

Leubner G, Ayele BT, Izydorczyk MS, Tuan PA, Sun M (2020) Ethylene regulates post-germination seedling growth in wheat through spatial and temporal modulation of ABA/GA balance. Journal of Experimental Botany 71: 1985–2004

Liu C, Jun JH, Dixon RA (2014) MYB5 and MYB14 Play Pivotal Roles in Seed Coat Polymer Biosynthesis in Medicago truncatula. Plant Physiology 165: 1424–1439

Liu M, Li Z, Zhang Y, Gao J (2020) Role of ethylene response factors (ERFs) in fruit ripening. Food Quality and Safety 4: 15–20

Ma W, Xu L, Gao S, Lyu X, Cao X, Yao Y (2021) Melatonin alters the secondary metabolite profile of grape berry skin by promoting VvMYB14-mediated ethylene biosynthesis. Horticulture Research 8

Mo R, Han G, Zhu Z, Essemine J, Dong Z, Li Y, Deng W, Qu M, Zhang C, Yu C (2022) The Ethylene Response Factor ERF5 Regulates Anthocyanin Biosynthesis in ‘Zijin’ Mulberry Fruits by Interacting with MYBA and F3H Genes. International Journal of Molecular Sciences 23

Mu H, Li Y, Yuan L, Jiang J, Wei Y, Duan W, Fan P, Li S, Liang Z, Wang L (2023) MYB30 and MYB14 form a repressor–activator module with WRKY8 that controls stilbene biosynthesis in grapevine. The Plant Cell 35: 552–573

Pan L, Zheng J, Liu J, Guo J, Liu F, Liu L, Wan H (2019) Analysis of the ASMT Gene Family in Pepper (Capsicum annuum L.): Identification, Phylogeny, and Expression Profiles. International Journal of Genomics 2019: 1–11

Pan Y, Seymour GB, Lu C, Hu Z, Chen X, Chen G (2011) An ethylene response factor (ERF5) promoting adaptation to drought and salt tolerance in tomato. Plant Cell Reports 31: 349–360

Pirrello J, Prasad BCN, Zhang WS, Chen KS, Mila I, Zouine M, Latché A, Pech JC, Ohme-Takagi M, Regad F, Bouzayen M (2012) Functional analysis and binding affinity of tomato ethylene response factors provide insight on the molecular bases of plant differential responses to ethylene. Bmc Plant Biology 12

Priest HD, Filichkin SA, Mockler TC (2009) cis-Regulatory elements in plant cell signaling. Current Opinion in Plant Biology 12: 643–649

Roldan MB, Cousins G, Fraser K, Hancock KR, Collette V, Demmer J, Woodfield DR, Caradus JR, Jones C, Voisey CR (2019) Elevation of Condensed Tannins in the Leaves of Ta-MYB14-1 White Clover (Trifolium repens L.) Outcrossed with High Anthocyanin Lines. Journal of Agricultural and Food Chemistry 68: 2927–2939

Serradilla MJ, Falagán N, Bohmer B, Terry LA, Alamar MC (2019) The role of ethylene and 1-MCP in early-season sweet cherry ‘Burlat’ storage life. Scientia Horticulturae 258

Sharafi Y, Jannatizadeh A, Fard JR, Aghdam MS (2021) Melatonin treatment delays senescence and improves antioxidant potential of sweet cherry fruits during cold storage. Scientia Horticulturae 288

Shelton D, Stranne M, Mikkelsen L, Pakseresht N, Welham T, Hiraka H, Tabata S, Sato S, Paquette S, Wang TL, Martin C, Bailey P (2012) Transcription Factors of Lotus: Regulation of Isoflavonoid Biosynthesis Requires Coordinated Changes in Transcription Factor Activity. Plant Physiology 159: 531–547

Sreekumar S, Sithul H, Muraleedharan P, Azeez JM, Sreeharshan S (2014) Pomegranate Fruit as a Rich Source of Biologically Active Compounds. BioMed Research International 2014: 1–12

Sun C, Liu L, Wang L, Li B, Jin C, Lin X (2020) Melatonin: A master regulator of plant development and stress responses. Journal of Integrative Plant Biology 63: 126–145

Sun LA, Zhang M, Ren J, Qi JX, Zhang GJ, Leng P (2010) Reciprocity between abscisic acid and ethylene at the onset of berry ripening and after harvest. Bmc Plant Biology 10

Tallapally M, Sadiq AS, Mehtab V, Chilakala S, Vemula M, Chenna S, Upadhyayula V (2020) GC-MS based targeted metabolomics approach for studying the variations of phenolic metabolites in artificially ripened banana fruits. Lwt 130

Wang J, Zhang H, Hou J, Yang E, Zhao L, Zhou Y, Ma W, Ma D, Li J (2023) Metabolic Profiling and Molecular Mechanisms Underlying Melatonin-Induced Secondary Metabolism of Postharvest Goji Berry (Lycium barbarum L.). Foods 12

Wang P, Yu A, Ji X, Mu Q, Salman Haider M, Wei R, Leng X, Fang J (2022) Transcriptome and metabolite integrated analysis reveals that exogenous ethylene controls berry ripening processes in grapevine. Food Research International 155

Wu K, Yu Y, Ni Y, Qiao T, Ji X, Xu J, Li B, Sun Q (2022) Overexpression of VvASMT1 from grapevine enhanced salt and osmotic stress tolerance in Nicotiana benthamiana. Plos One 17

Xu L, Xiang G, Sun Q, Ni Y, Jin Z, Gao S, Yao Y (2019) Melatonin enhances salt tolerance by promoting MYB108A-mediated ethylene biosynthesis in grapevines. Horticulture Research 6

Xu L, Yue Q, Bian Fe, Sun H, Zhai H, Yao Y (2017) Melatonin Enhances Phenolics Accumulation Partially via Ethylene Signaling and Resulted in High Antioxidant Capacity in Grape Berries. Frontiers in Plant Science 8

Xu L, Yue Q, Xiang G, Bian Fe, Yao Y (2018) Melatonin promotes ripening of grape berry via increasing the levels of ABA, H2O2, and particularly ethylene. Horticulture Research 5

Yang Q, Peng Z, Ma W, Zhang S, Hou S, Wei J, Dong S, Yu X, Song Y, Gao W, Rengel Z, Huang L, Cui X, Chen Q (2021) Melatonin functions in priming of stomatal immunity inPanax notoginseng and Arabidopsis thaliana. Plant Physiology 187: 2837–2851

Yilmaz Y, Göksel Z, Erdoğan SS, Öztürk A, Atak A, Özer C (2015) Antioxidant Activity and Phenolic Content of Seed, Skin and Pulp Parts of 22 Grape (Vitis vinifera L.) Cultivars (4 Common and 18 Registered or Candidate for Registration). Journal of Food Processing and Preservation 39: 1682–1691

Yu G, Wang L-G, He Q-Y (2015) ChIPseeker: an R/Bioconductor package for ChIP peak annotation, comparison and visualization. Bioinformatics 31: 2382–2383

Yue P, Wang Y, Bu H, Li X, Yuan H, Wang A (2019) Ethylene promotes IAA reduction through PuERFs-activated PuGH3.1 during fruit ripening in pear (Pyrus ussuriensis). Postharvest Biology and Technology 157

Zhang X, Abrahan C, Colquhoun TA, Liu C-J (2017) A Proteolytic Regulator Controlling Chalcone Synthase Stability and Flavonoid Biosynthesis in Arabidopsis. The Plant Cell 29: 1157–1174

Zhang X, Ma W, Guan X, Wang F, Fan Z, Gao S, Yao Y (2023) VvMYB14 participates in melatonin-induced proanthocyanidin biosynthesis by upregulating expression of VvMYBPA1 and VvMYBPA2 in grape seeds. Horticulture Research 10

